# A simple secretion assay for assessing new and existing myocilin variants

**DOI:** 10.1101/2021.10.30.466573

**Authors:** Emi Nakahara, John D. Hulleman

## Abstract

With the increasing use of molecular genetics approaches for determination of potential disease-causing mutations, it is becoming more important to be able to interpret and act upon the provided results. As an example of such an instance, nearly 300 mutations have been identified in the myocilin (*MYOC*) gene, which is the most commonly mutated gene causing primary open angle glaucoma. Yet a lack of sufficient information exists for many of these variants, hindering their definitive classification. While the function of MYOC is unclear, biochemically, the vast majority of glaucoma-causing MYOC mutations result in protein non-secretion and intracellular insoluble aggregate formation in cultured cells. Previously we generated a *Gaussia* luciferase-based MYOC fusion protein to sensitively track secretion of the protein. Herein we applied this same assay to fourteen clinically-derived MYOC variants with varying degrees of predicted pathogenicity and compared the luciferase secretion results with the better established MYOC assay of western blotting. Eight of the variants (G12R, V53A, T204T, P254L, T325T, D380H, D395_E396insDP, and P481S) had not been biochemically assessed previously. Of these, P254L and D395_E396insDP demonstrated significant secretion defects from human embryonic kidney (HEK-293A) cells reminiscent of glaucoma-causing mutations. Overall, we found that the luciferase assay results agreed with western blotting for thirteen of the fourteen variants (93%), suggesting a strong concordance. These results suggest that the *Gaussia* luciferase assay may be used as a complementary or standalone assay for quickly assessing MYOC variant behavior, and anticipate that these results will be useful in MYOC variant curation and reclassification.

## INTRODUCTION

Glaucoma is the leading cause of irreversible blindness worldwide, accounting for 11% of the total cases of vision loss^1^. Primary open angle glaucoma (POAG), the most common subset of glaucoma, is a chronic, ocular hypertensive disease that ultimately triggers the death of retinal ganglion cells. The myocilin (*MYOC*) gene was the first gene to be associated with POAG, and its mutation is also the most common cause of glaucoma^2^. While glaucoma-associated mutations in *MYOC* were identified nearly 25 years ago^3^, little is still known regarding the normal function of the MYOC protein. Wild-type MYOC is a secreted, extracellular matrix protein that is expressed throughout the eye, including in the trabecular meshwork, which is a key tissue involved in regulating aqueous outflow and governance of intraocular pressure (IOP)^4^. Glaucoma-causing autosomal dominant mutations in MYOC lead to protein aggregation^5-7^, are poorly secreted^7-10^, linked to IOP elevation^11, 12^, and trabecular meshwork death/dysfunction^13, 14^.

Yet, unless there is a strong genetic correlation between the mutation and disease in multiple individuals, it is many times difficult to assess whether a variant is truly pathogenic, and therefore a possible therapeutic target. Accordingly, the American College of Medical Genetics (ACMG) and the Association for Molecular Pathology (AMP) have developed recommendations for improving the consistency of clinical variant interpretation^15^, one aspect of which is to use a “well-established” functional assay to provide support for either a benign or pathogenic effect^16^. This evidence can be used to provide information regarding the effect of the mutation on protein function and can even lead to the reclassification of certain variants^17^.

Previously, we have utilized the naturally secreted *Gaussia* luciferase (eGLuc2)^18^ as a means to conveniently monitor the secretion of engineered and disease-associated mutations in the fibulin-3 protein^19-21^, and more recently for evaluating the potential effects of mutations on MYOC secretion^8^. In the latter publication, we extensively compared the biochemical characteristics of WT and Y437H MYOC as FLAG and eGLuc2 fusions but stopped short of expanding this direct comparison to additional MYOC mutations. Therefore, while the eGLuc2 tag has no effect on MYOC secretion propensity, and the assay is inexpensive, easy to use, convenient, and potentially high throughput^22^, until now, it has not been thoroughly vetted as a true functional assay for a variety of MYOC mutations through direct comparison to conventional MYOC biochemical assays (protein non-secretion/intracellular insoluble aggregation determined by western blotting)^5-7, 9, 10, 14, 23^. Herein, we sought to characterize fourteen mutations in MYOC that had previously not been tested as MYOC eGLuc2 fusion proteins and compare/contrast these results with the equivalent mutations generated as FLAG-tagged MYOC proteins and quantified by western blotting. We found that the luciferase assay results agreed with western blotting for thirteen of the fourteen variants (93%), demonstrating a strong concordance. Our results suggest that luminescent analysis of MYOC eGLuc2 can be used for quickly assessing MYOC variant behavior.

## MATERIALS AND METHODS

### Plasmid Generation

The original pcDNA3 wild-type (WT) and Y437H MYOC plasmids with a C-terminal FLAG-tag (FT) were provided by Charles Searsby and Dr. Val Sheffield (University of Iowa). The Q5 Site-Directed Mutagenesis Kit (New England Biolabs (NEB), Ipswich, MA, USA) was used to create the fourteen different MYOC mutant constructs using the pcDNA3 WT MYOC DNA as a template. The enhanced *Gaussia* luciferase 2 (eGLuc2 [containing the M43I and M110I mutations that eliminate oxidation-prone methionines], also C-terminal) FT MYOC variants were generated by DNA assembly (NEBuilder HiFi DNA Assembly Cloning Kit, NEB) by combining a PCR-amplified eGLuc2 product from a pcDNA3 MYOC eGLuc2 plasmid (originally described previously^8^), with a PCR product containing the remainder of each pcDNA3 MYOC FT plasmid. The final assembled construct encoded MYOC followed by a 54 bp linker, the eGLuc2 tag and the FLAG tag. All constructs were verified through full Sanger sequencing.

### Cell Culture

Human embryonic kidney cells (HEK-293A, Life Technologies) were cultured in high glucose (4.5 g/L) Dulbecco’s modified Eagle’s Medium (DMEM; Corning, Corning, NY, USA) supplemented with 10% fetal bovine serum (Omega Scientific, Tarzana, CA, USA) and 1% penicillin/streptomycin/L-glutamine (Corning). Cells were grown at 37°C and 5% CO_2_.

### Transfection

For transfections with MYOC FT constructs, HEK-239A cells were plated in 12-well plates (Costar, Corning) at a concentration of 110,000 cells/well and allowed attachment overnight. For MYOC eGLuc2 FT transfections, cells were plated overnight at a density of 55,000 cells/well in a 24 well plate (Costar, Corning). The pcDNA3 MYOC FT and pcDNA3 MYOC eGLuc2 FT constructs were introduced into the HEK cells using Lipofectamine 3000 (Life Technologies, Carlsbad, CA, USA) at a concentration of 1 μg/well (3 μL of Lipofectamine) and 500 ng/well (1.5 μL Lipofectamine), respectively. A pEGFP-C1 (Takara, San Jose, CA, USA) construct served as a control to assess transfection efficiency. Twenty-four hours after transfection, media was then changed with low serum media (0.5-1% FBS) to minimize FBS-related disruptions in western blotting of the conditioned media. Cells and media were then collected 24 h later, 48 h post initial transfection and subsequently used for either western blotting or a luciferase assay.

### Gaussia luciferase assay

For the secreted fraction samples, fifty uL aliquots of conditioned media were collected 24 h following media change (48 h post transfection). Media was transferred to a 96-well polystyrene flat-bottomed black assay microplate (Costar, Corning). One hundred nL of 50x coelenterazine substrate diluted in 10 μL of NanoFuel® GLOW reagent for Gaussia Luciferase (NanoLight Technology, Prolume, Pinetop, AZ) was then added to the media samples. Following 5 min of incubation at RT in the dark, the luminescence was measured immediately with a BioTek Synergy 2 plate reader (BioTek, Winooski, VT, USA) using a 1 sec integration time. For the luciferase assay, * = p < 0.05, ** = p < 0.01, *** = p < 0.001 using a 1 sample t-test against a hypothetical value of 1 (i.e., unchanged relative to WT). n ≥ 5 independent experiments.

### Preparation of Insoluble MYOC

Cells transfected with MYOC FT constructs were lysed in well using PBS (Corning) supplemented with 0.1% Triton X-100 (Fisher, Waltham, MA, USA) and Halt protease inhibitors (Pierce Thermo Fisher, Rockford, IL, USA) followed by rocking for 30 min at 4°C. The insoluble fraction was pelleted by centrifugation at 14,800 x *g* for 10 minutes at 4°C and washed once in cold PBS (followed by centrifugation). The remaining soluble protein fraction was collected and quantified for protein concentration via bicinchoninic acid (BCA) assay (Pierce Thermo Fisher). The soluble protein concentration was then used as an indication of lysis efficiency/cell mass to determine the amount of 8 M urea (dissolved in PBS, Fisher) used to resuspend the insoluble protein pellet.

### Western Blotting

Media (40 μL) and insoluble samples and were denatured in reducing Laemmli buffer at 100°C for 5-10 min prior to loading onto a 4-20%Tris-Gly SDS-PAGE gel (Life Technologies). Proteins were then transferred to a nitrocellulose membrane using an iBlot2 (Life Technologies). Ponceau S (Sigma, St. Louis, MO) reversible staining of the blots qualitatively verified equal loading across samples. Blots were blocked overnight at 4°C in PBS Odyssey Blocking Buffer (LI-COR, Lincoln, NE, USA), and then probed with either a goat anti-MYOC antibody (cat# sc-21243 [no longer made], 1:1000; Santa Cruz, Dallas, TX), or a rabbit anti-FLAG antibody (cat# PA1-984B, 1:1000; Thermo Fisher Scientific) for 1 h at RT followed by an appropriate near-infrared secondary antibody (1:15,000; LI-COR). A LI-COR Odyssey CLx infrared scanner (LI-COR) was used to image blots, and Image Studio software (LI-COR) was used to quantify protein bands. For western blotting, * = p < 0.05, ** = p < 0.01, *** = p < 0.001 using a 1 sample t-test against a hypothetical value of 1 (i.e., unchanged relative to WT for secreted) or a 2 sample t-test compared to WT MYOC for insoluble protein. n ≥ 4 independent experiments.

## RESULTS

Glaucoma-related mutations in MYOC have been demonstrated to cause protein misfolding, non-secretion or inefficient secretion, and an increase in Triton-X insoluble protein^5-8, 10, 14, 23^. Accordingly, many pathogenic MYOC mutations have also been demonstrated to increase levels of endoplasmic reticulum (ER) stress^11, 12, 24-26^. Yet whether these same mutations cause higher IOP and hence glaucoma directly through an ER stress mechanism is still under investigation. Nonetheless it is clear that potentially pathogenic MYOC variants can be distinguished from benign mutants through their biochemical characteristics in an immortalized cell culture system. In this work, we asked whether a simple *Gaussia* luciferase-based MYOC secretion assay could be used to complement or replace the more cumbersome western blotting approach for MYOC variants.

HEK-293A cells were transfected with sixteen total constructs (Fig. 1A), including a WT MYOC eGLuc2 FT control and an established pathogenic control, Y437H MYOC eGLuc2 FT. The fourteen remaining variants (G12R, Q48H, V53A, R76K, T204T, P254L, T325T, G367R, D380H, D395_E396insDP, E396dup, A445V, N480K, and P481S) were selected because they range in predicted behavior from ‘benign’ to ‘variant of uncertain significance (VUS)’ to ‘pathogenic’ according to ClinVar (https://www.ncbi.nlm.nih.gov/clinvar/) and a MYOC mutation database (www.myocilin.com). Whereas the majority of the mutations that we analyzed were missense mutations, to be more encompassing of the variants identified in the *MYOC* gene, we also included two synonymous mutations, T204T (c.855G>T, identified in one POAG patient and no controls) and T325T (c.975G>A, identified in no POAG patients, and two controls)^27^. Importantly, within the set of fourteen variants, we also purposely included five mutants (Q48H, R76K, G367R, E396dup, and A445V) that had been characterized previously in publications to serve as calibrators for our assay. Importantly, each of these MYOC eGLuc2 FT ‘calibrators’ displayed secretion profiles in accordance with past results (Fig. 1B), namely WT-like secretion of Q48H, R76K, and A445V and poor secretion of G367R and E396dup (Fig. 1B). These results demonstrate an alignment of observations that was consistent across two different groups and using different biochemical approaches^10^. Of the remaining previously uncharacterized MYOC variants (G12R, V53A, T204T, P254L, T325T, D380H, D395_E396insDP, and P481S), only P254L and D395_E396insDP demonstrated significant secretion defects using the *Gaussia* luciferase assay (Fig. 1B), hinting that these mutations are potentially pathogenic.

**Figure 1.**
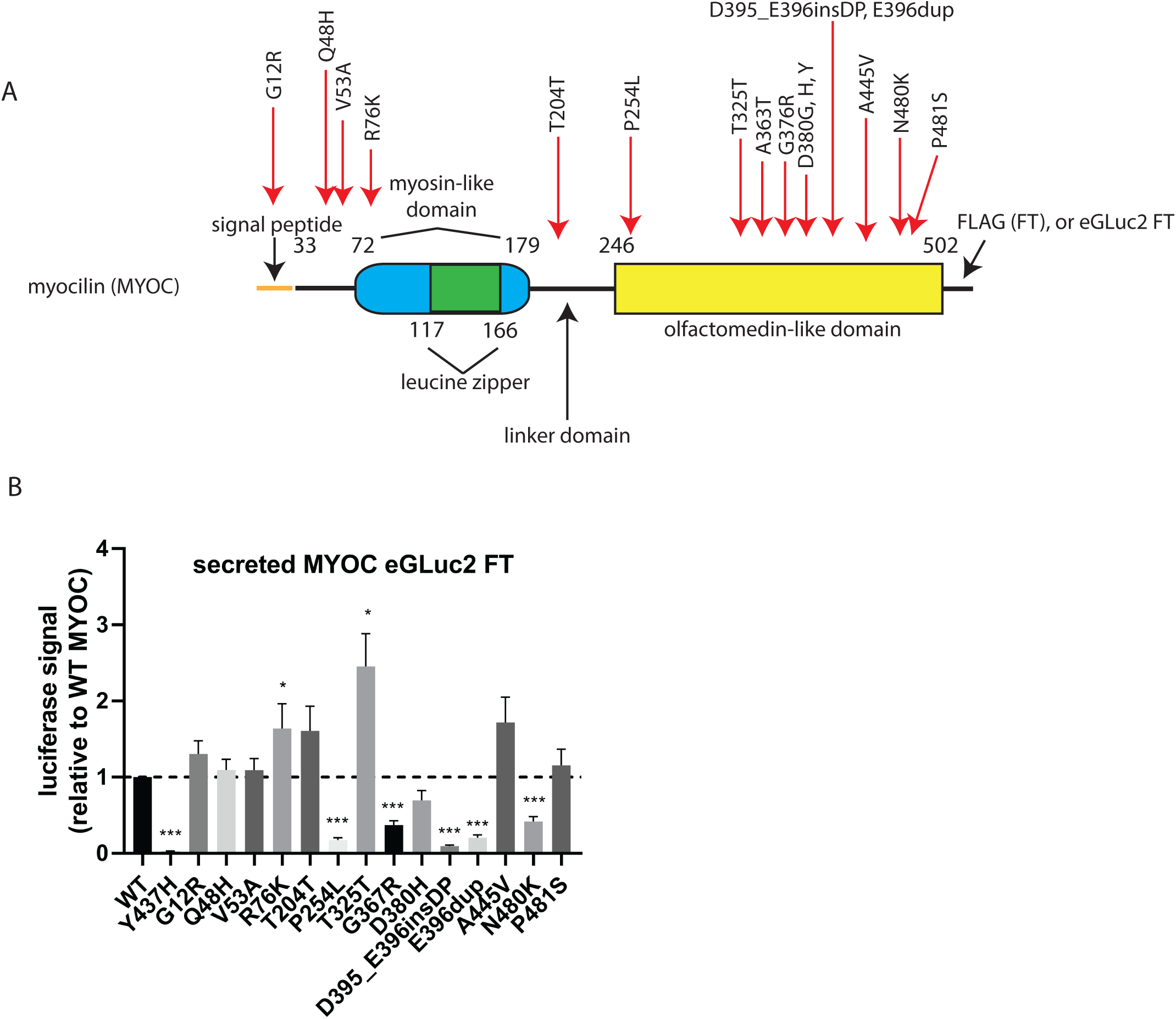
Screening of fourteen MYOC eGLuc2 FT variants demonstrates a range of secretion characteristics. (A) Cartoon schematic of MYOC and the generated mutations. (B) Secreted *Gaussia* luciferase (eGLuc2) luminescence results after transfection of the indicated constructs in HEK-293A. * = p < 0.05, ** = p < 0.01, *** = p < 0.001 using a 1 sample t-test against a hypothetical value of 1 (i.e., unchanged relative to WT). n ≥ 5 independent experiments.

In parallel to the above-described *Gaussia* luciferase-based detection of MYOC secretion, we also performed western blotting on secreted (Fig. 2A, B) and insoluble (Fig. 2A, C) MYOC FT (not eGLuc2-tagged) to more closely mimic the conventional biochemical characterization of MYOC variants by western blotting^5, 6, 9, 10, 14, 28^. Interestingly, when comparing secreted MYOC, the *Gaussia* luciferase assay agreed with western blotting for thirteen of the fourteen variants tested (93%), excluding D380H (cf. Fig. 1B to Fig. 2B). MYOC FT variants that demonstrated significant secretion defects (P254L, G367R, D380H, D395_E396insDP, E396dup, and N480K) were all found to have significantly higher insoluble intracellular levels (Fig. 2A, C). While the secreted data for nearly all variants were quite consistent across assays, the P481S variant seemed to have a unique behavior of being secreted efficiently, but also having significantly higher insoluble levels as detected by western blotting (Fig. 2A-C). This behavior of partial solubility has also been demonstrated with other MYOC variants including G364V, T377M, D380A, and R422C^5^. Such an observation will likely complicate applying functional rules to determine whether its behavior supports benign or pathogenic classification.

**Figure 2.**
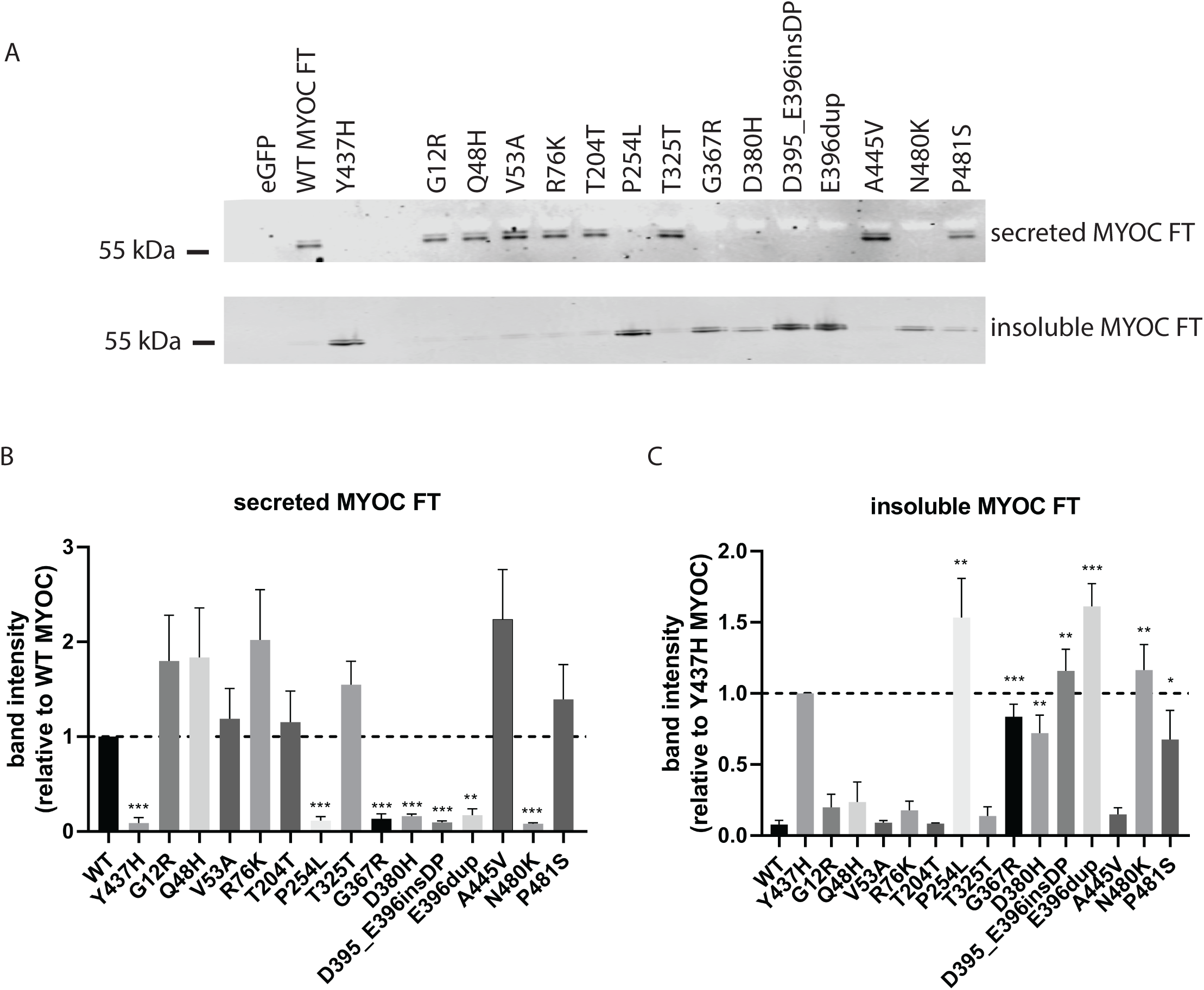
Parallel analysis of MYOC variants by western blotting demonstrates a strong concordance with the MYOC eGLuc2 FT results. (A) Western blotting results of secreted and insoluble intracellular levels of transfected MYOC FT constructs. (B, C) Quantification of secreted (B) and insoluble (C) MYOC FT levels from western blotting performed in (A). Because of the more reliable signal obtained from insoluble Y437H MYOC FT, insoluble levels of new variants were first compared to those of Y437H MYOC FT, and then we performed a significance test asking whether this obtained value was significantly different from WT MYOC FT insoluble levels. * = p < 0.05, ** = p < 0.01, *** = p < 0.001 using a 1 sample t-test against a hypothetical value of 1 (i.e., unchanged relative to WT MYOC for secreted) or a 2 sample t-test compared to WT MYOC for insoluble protein. n ≥ 4 independent experiments.

## CONCLUSIONS/DISCUSSION

In summary, we found that the assay results from our *Gaussia* luciferase fusion constructs agreed with secretion data from western blotting for thirteen of the fourteen variants (93%), suggesting a strong concordance, and increase the likelihood that this assay could be used as a surrogate, or in addition to conventional western blotting for MYOC. For example, the luciferase assay may be used as an initial screening method to identify possibly pathogenic variants from a large set of mutant MYOCs, followed by confirmation of the behavior of those variants via western blotting. While these studies were mainly performed to directly compare the luciferase assay to western blotting, they also reveal some notable points. The first point highlights the potential importance of functional evidence for variant reclassification^16^, which is especially important given the lack of such evidence for most MYOC variants and that around half of all variants in ClinVar are currently classified as VUS. The second point is that we have demonstrated for the first time that the P254L^29^ and D395_E396insDP^30^ mutations appear to behave in a manner consistent with pathogenic MYOC. The third point is that our results would suggest that the V53A, Q48H, and R76K variants are likely benign since they are readily secreted and soluble. These observations would potentially clarify their ClinVar classification of “uncertain significance: and “conflicting interpretations of pathogenicity”, respectively. Fourthly, the discrepancy between the *Gaussia* luciferase fusion and FT version of D380H highlights the potentially difficulty with applying one single criteria or assay to help in reclassification strategies based on function. This residue appears to be a hot spot mutation involved in calcium binding; additional D380Y, D380G and D380A mutations have been observed in the population^31, 32^. Intriguingly, a recent study has shown that mutation of D380 can result in a stable, but non-native structure compared to the WT MYOC olfactomedin domain^33^. Moreover, the D380A mutation was found to have ‘partial solubility’ in previous studies, indicating a unique intermediate behavior between what would be considered benign vs. pathogenic. Overall, these observations indicate the potential challenges in classification of certain MYOC variants that do not seem to abide by more common MYOC behaviors. Yet, in conclusion, our results suggest that the *Gaussia* luciferase assay maybe used as a complementary or standalone assay for quickly assessing MYOC variant behavior, and we predict that these observations will be useful in MYOC variant curation and reclassification.

## ACKNOWLEDGEMENTS

The work described herein was supported by an endowment from the Roger and Dorothy Hirl Research Fund and a National Eye Institute Visual Science Core Grant (P30 EY030413).

